# Representation of conscious percept without report in the macaque face patch network

**DOI:** 10.1101/2020.04.22.047522

**Authors:** Janis K. Hesse, Doris Y. Tsao

**Author notes:** Corresponding authors: Janis K. Hesse, California Institute of Technology, Computation and Neural Systems, MC 136-93 Pasadena, CA 91125, USA, Doris Y. Tsao, Professor of Biology Investigator, HHMI, California Institute of Technology, Division of Biology, MC 114-96 Pasadena, CA 91125, USA.

## Abstract

A powerful paradigm to identify the neural correlates of consciousness is binocular rivalry, wherein a constant visual stimulus evokes a varying conscious percept. It has recently been suggested that activity modulations observed during rivalry could represent the act of report rather than the conscious percept itself. Here, we performed single-unit recordings from face patches in macaque inferotemporal (IT) cortex using a no-report paradigm in which the animal’s conscious percept was inferred from eye movements. We found high proportions of IT neurons represented the conscious percept even without active report. Population activity in single trials, measured using a new 128-site Neuropixels-like electrode, was more weakly modulated by rivalry than by physical stimulus transitions, but nevertheless allowed decoding of the changing conscious percept. These findings suggest that macaque face patches encode both the physical stimulus and the animal’s conscious visual percept, and the latter encoding does not require active report.

## Introduction

Having conscious experience is arguably the most important reason why it matters to us whether we are alive or dead. The question which signals in the brain reflect this conscious experience and which reflect obligatory processing of input regardless of conscious experience is therefore one of the most important puzzles in neuroscience. For example, activations in the retina may correlate with the conscious percept of flashing light but are arguably entirely driven by physical input, much of which never evolves into a conscious percept. Another driver of neural activity that can be confounded with signals related to conscious perception is report. Recently, it has been suggested that brain regions may correlate with conscious perception simply because they are driven by the active report of it (Frässle et al., 2014; Koch et al., 2016; Overgaard & Fazekas, 2016; Tsuchiya et al., 2015).

A paradigm known as binocular rivalry is useful for distinguishing responses related to conscious perception from those driven by obligatory processing of physical input (Blake et al., 2014; Tong et al., 2006): When two incompatible stimuli such as a face and an object are shown to the left and right eyes, respectively, one does not perceive a constant superimposition of the two, but instead one’s percept alternates between face and object even though the physical input is fixed (Fig. 1a). Since these alternations are internally generated, they cannot be attributed to pure feedforward processing of external input.

**Figure 1.**
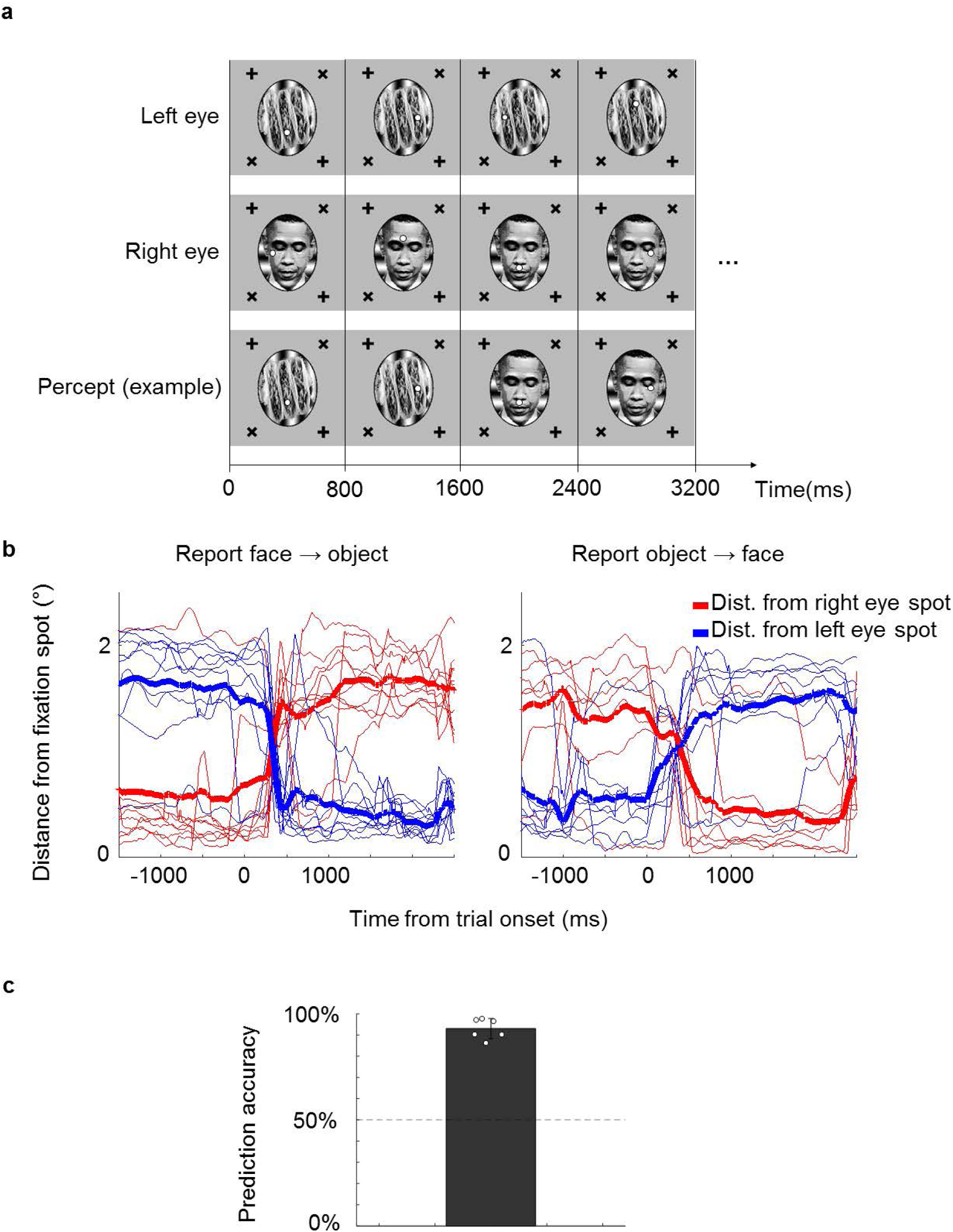
A novel no-report paradigm. **(a)** Illustration of binocular rivalry stimuli used in the paradigm. Four example trials are shown. Each trial was presented continuously for 800 ms each without blank period between trials. The first and second row show stimuli in the left and right eyes, respectively. If different stimuli are shown to the left and right eye, as in this example, one’s percept will spontaneously alternate between the two, as shown in the example perceptual trajectory in the third row. Stimuli in each eye contained a fixation spot at one of four possible positions that the monkey was trained to fixate on. **(b)** Example eye traces from a human subject. Red and blue traces show the distance of the eye position from the fixation spot that is shown in the right and left eye, respectively. Thick lines show the average. Traces are aligned to the onset of a trial where the subject reported that the percept switched from face to object (left), or object to face (right). **(c)** The bar plot shows the average proportion of those trials where the percept inferred matched the percept reported by button press. White circles show accuracies of individual subjects. We inferred that a subject was perceiving face or object if the subject fixated on the face fixation spot (i.e., fixation spot in the eye of the face stimulus) or object fixation spot (i.e., fixation spot in the eye of the object stimulus), respectively, for at least half of the trial.

In previous studies, researchers trained monkeys to report their percept during binocular rivalry by releasing a lever and found that the proportion of cells modulated by the reported percept increases along the visual hierarchy, with as little as 20% of cells showing modulations in V1 (Leopold & Logothetis, 1996) compared to 90% of cells showing modulations in IT (Sheinberg & Logothetis, 1997). Using fMRI, Tong et al. found that the human fusiform face area responds to reported perceptual switches (Tong et al., 1998). The reported percept also modulates activity of single units in the human medial temporal lobe and frontal cortex (Gelbard-Sagiv et al., 2018).

Although binocular rivalry isolates the conscious percept from physical input, an important confounding factor remains. In all studies cited above, the monkey or human subject always actively reported their percept by a motor response. Thus it is possible that the observed neuronal activations were due to the act of report itself, including introspection, decision making, and motor action accompanying report, rather than a switch in conscious percept (Koch et al., 2016; Overgaard & Fazekas, 2016; Tsuchiya et al., 2015). This concern was emphasized in an fMRI experiment by Frässle et al. who compared modulations in the brain with and without active report (Frässle et al., 2014). Many of the modulations observed in higher-level brain regions such as the frontal lobe disappeared when subjects did not actively report perceptual switches.

To infer the subject’s percept in the absence of report, Frässle et al. used two no-report paradigms that depended on pupil size and optokinetic nystagmus, respectively. If the stimuli in the two eyes have different brightness, the subject’s pupil size will vary according to the dominant percept’s brightness and can thus be used to infer the percept. As a second method, Frässle et al. exploited optokinetic nystagmus. They presented gratings moving in opposite directions in the two eyes, causing the subject’s eye position to reflexively follow the direction of the dominant grating.

These no-report paradigms allow accurate prediction of the subject’s percept but are not free of confounds themselves (Overgaard & Fazekas, 2016). First, pupil size is known to correlate with arousal, surprise, attention and other confounding factors (Bradley et al., 2008; Hoeks & Levelt, 1993; Preuschoff et al., 2011). Second, when optokinetic nystagmus is applied to moving non-grating stimuli such as natural objects that drive IT cortex, there will be confounding physical stimulus differences. For example, the dominant stimulus that is smoothly pursued by the subject’s eyes will tend to be stationary on the subject’s fovea and optimally modulate IT areas with foveal biases, while the non-dominant stimulus will be more eccentric and have increased motion velocity. Moreover, optokinetic nystagmus is still present in monkeys where the conscious percept is diminished due to anesthesia with low doses of ketamine (Leopold et al., 2002).

Here, we introduce a new no-report paradigm that relies on active tracking of a fixation spot, unlike the reflex-based paradigms mentioned above. In this fixation-based paradigm the subject is required to maintain fixation on a jumping spot, a task that many animals in vision research are already trained to perform. While following the fixation spot, subjects view either unambiguous, monocular stimuli physically switching between a face and an object, or a binocular rivalry stimulus that switches only perceptually. For the binocular rivalry stimulus, a fixation spot is shown to each eye at different positions on the screen. Thus, when the subject perceives a face in the left eye, he/she will generally perceive only the fixation spot in the left eye and saccade to it, ignoring the fixation spot in the right eye. In this way, the subject’s percept can be inferred from his/her eye movement patterns without active report.

In a second innovation, we performed electrophysiological recordings using a novel 128-electrode site Neuropixels-like probe that allowed us to measure responses from large numbers of cells simultaneously. This allowed us to address for the first time the extent to which neural activity is modulated by conscious perception *in single trials*. Sheinberg and Logothetis (1997) found that 90% of IT cells were modulated by conscious perception, but the response modulations reported in that study during the rivalry condition were clearly smaller than those in the physical condition. This decrease could have been due to mixed selectivity of cells for the conscious percept and the physical stimulus on single trials. Alternatively, cells could have been modulated just as strongly by perceptual as by physical alternations and the decrease could have been due to incorrect reporting of the percept on some trials. Inter-trial averaging confounds these two possibilities.

To explore correlates of conscious perception, we targeted recordings to macaque face patches ML and AM. The macaque face patch system constitutes an anatomically connected network of regions in IT cortex dedicated to face processing (Chang & Tsao, 2017; Grimaldi et al., 2016; Tsao et al., 2006). To date, most response properties of cells in the face patch network can be explained in a feedforward framework without invoking conscious perception. For example, the functional hierarchy of this network, with increasing view invariance as one moves anterior from ML to AM (Freiwald & Tsao, 2010), can be explained by simple feedforward pooling mechanisms (Leibo et al., 2017). The representation of facial identity by cells in face patches through projection onto specific preferred axes can also be explained by feedforward mechanisms (Chang & Tsao, 2017). At the same time, it has been postulated that the fundamental architecture of the cortex may be a predictive loop, in which inference guided by internal priors plays a key role in determining what we see (Rao & Ballard, 1999). For example, one explanation for binocular rivalry is that it directly reflects our knowledge that two objects can’t occupy the same space (Hohwy et al., 2008). The hierarchical organization of the face patch network, together with its specialization for a single visual form, make it a promising testbed to examine the neural circuits underlying construction of conscious visual experience, beyond feedforward filtering of visual input.

Here, we recorded from fMRI-identified face patches ML and AM in two monkeys using high-channel electrodes while we inferred the animals’ conscious percept through the no-report paradigm described above. We found that high proportions of cells in both face patches (61% in ML and 81% in AM) encode the conscious percept even without active report. Population activity of perceptually-modulated cells was more weakly modulated during rivalry than during physical stimulus transitions in single trials. Nevertheless, we could still reliably decode the dynamically changing percept. Overall, these findings suggest that cells in macaque face patches encode both the physical stimulus and the animal’s conscious visual percept.

## Results

We first confirmed that it is possible to correctly infer a subject’s conscious percept using a fixation-based no-report paradigm through a behavioral experiment in humans. We presented binocular rivalry stimuli consisting of a face (e.g., Obama) in the right eye and a non-face object (e.g., a taco) in the left eye, causing the percept to stochastically alternate between the two (Fig. 1a). Each of the stimuli contained a fixation spot that jumped to one of four possible locations every trial. Trials were 800 ms long and contained no blank period, i.e., stimuli were presented continuously. If subjects fixated at the fixation spot presented in the right eye on a given trial, we inferred that they perceived the face and vice versa for the object. To verify that the percept of face or object could be inferred from fixations, we instructed 6 naïve human subjects to perform the fixation task while simultaneously reporting their conscious percept with button presses. On trials where the percept switched, subjects also switched the fixation spot they were following (Fig. 1b). We were able to infer which image the subjects were consciously perceiving with accuracies ranging from 86% to 98% across subjects (average: 93%, Fig. 1c).

We next used the same method in monkeys to infer their conscious percept while recording from face patches ML and AM in IT. Importantly, the two monkeys in this study had never been trained to report their percept. They had previously been trained to maintain fixation on a spot (presented binocularly) so they learned to perform the task within one or two days, respectively (reaching performance of maintaining fixation on a spot on at least 80% of all trials). We presented two types of stimuli: In the “physical” condition, unambiguous monocular stimuli were physically switched between face and object. In the “perceptual” (binocular rivalry) condition the same face and object were continuously presented to the right and left eye, respectively, so any changes in percept were internally generated. To account for individuals’ eye dominance, we balanced the contrasts of the stimuli in the two eyes so that the monkey followed both fixation spots equally often in the rivalry condition. We inferred switches during rivalry when monkeys behaviorally switched from following the fixation spot in one eye to following the fixation spot in the other eye, as shown in the example eye traces in Fig. 2a, top. Spike rasters aligned to onset of trials where the percept switched from an example ML cell recorded in the same session are shown in Fig. 2a, bottom. Fig. 2b compares average response time courses to physical switches to face or object with responses to perceptual switches in example cells from ML and AM. Both example cells responded more strongly to a physically presented face than object, which is expected since they were recorded from face patches. Importantly, in the binocular rivalry condition, when the monkey perceived a face as inferred by its eye movement, the response of both cells was also higher than when the monkey perceived an object. Since the physical stimulus was identical in both cases, the response reflected its conscious percept of a face rather than just the physical input.

**Figure 2.**
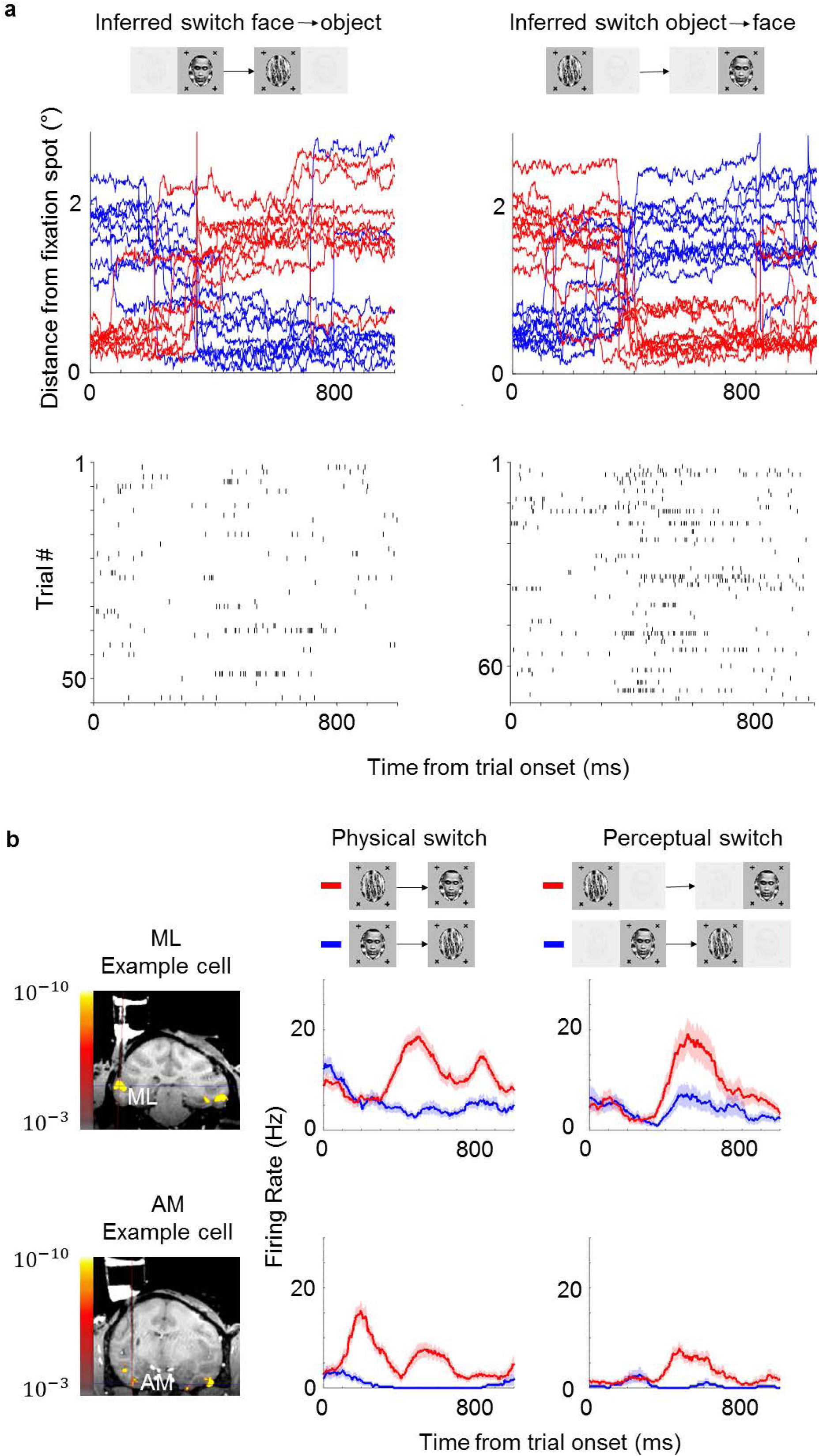
Example face cells modulated by both physical and perceptual switches to face. **(a)** Top: Example eye traces from a macaque performing the task aligned to a trial where the inferred percept switched from face to object (left) and from object to face (right), respectively. Red and blue curves indicate distances from the face and object fixation spots, respectively (as in Fig. 1b). Bottom: Spike raster of an example ML cell recorded in the same session as for the top panel. Responses are aligned to all trials where the inferred percept switched from face to object (left) and from object to face (right), respectively. **(b)** Left: Coronal slices from magnetic resonance imaging scan showing recording locations for the two example cells in this figure (top: face patch ML, bottom: face patch AM). Color overlay shows functional MRI activation to visually presented faces vs non-face objects. Middle: Peristimulus histograms (PSTHs) show neuronal response time courses aligned to trial onsets where the visual stimulus was physically switched from face to object (blue) or from object to face (red). Right: PSTHs aligned to trial onsets where the inferred percept switched from face to object (blue) or object to face (red). ML cell is same cell as in (a). Shaded areas indicate standard error mean across trials.

We recorded a total of 347 cells in ML and 210 cells in AM that were selective, i.e., showed a significant difference between face and object in the physical switch condition (p<0.05, two-sided t-test). Population results of all selective cells are shown in Fig. 3. Since we recorded from face patches, most cells showed stronger responses to the physically presented face stimulus. Importantly, most cells kept their preference in the perceptual condition. In face patch ML, 61% of cells were significantly modulated by the conscious percept in the binocular rivalry condition and showed preference consistent with the physical switch condition (p<0.05, two-sided t-test), while 9% of cells were significantly but inconsistently modulated. In AM, a face patch that receives input from ML (Grimaldi et al., 2016) and is the highest patch in the face patch hierarchy within IT (Freiwald & Tsao, 2010), the percentage of consistent modulation increased to 81%, with only 1% showing inconsistent modulation. For both patches there was a clear correlation between modulation by physical stimuli and modulation by the percept in binocular rivalry (*r* = *0*.*72, p* < 10^−31^). Thus, in a no-report paradigm, cells in IT exhibit modulations by the conscious percept that reflect their response tuning to physically unambiguous inputs.

**Figure 3.**
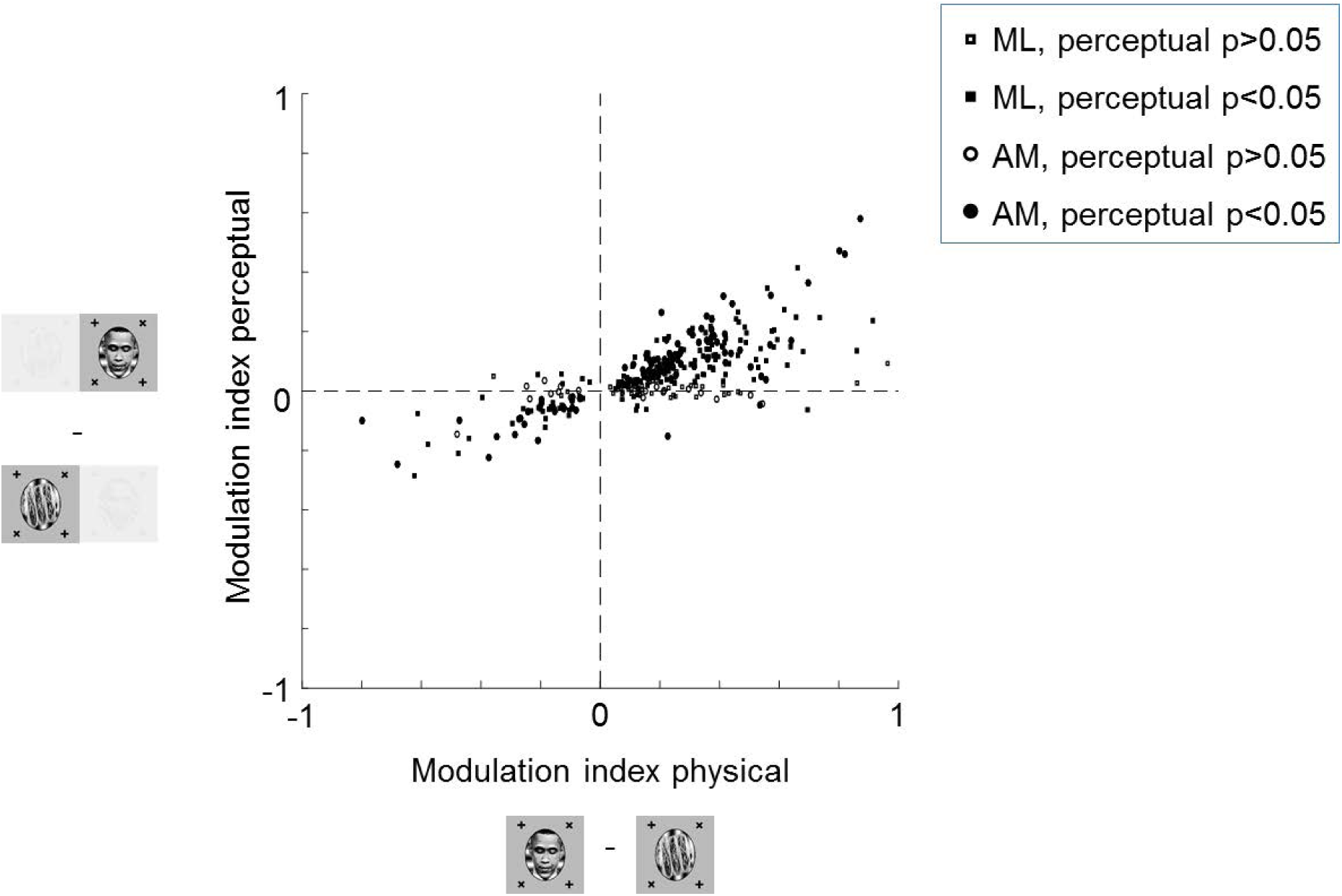
High proportions of face cells show modulation by conscious percept. Scatter plot shows modulation indices 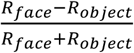 measuring the difference in responses (i.e. average spike count *R*) on trials where the inferred percept was face or object, respectively, for the physical monocular condition (x-axis) and perceptual binocular rivalry condition (y-axis). Squares show cells from ML and circles show cells from AM. Open and filled markers indicate cells without and with significant difference between perceived face and perceived object response in the binocular rivalry condition, respectively.

After eliminating the report confound, two important potential confounds remain: First, cells could be selective for the eye-of-origin of the fixation point that the animal is following (e.g., a cell could respond selectively to a fixation spot in the fovea of the left eye). Second, since we presented binocular stimuli using red-cyan anaglyph goggles, a confound could arise if cells were selective for the color of the fixation spot that is in the fovea. To control for these two potential confounds, we switched the colors and eye-of-origin of the face and object stimuli, i.e., where the face and its corresponding fixation spot was previously presented in red in one eye, it was now presented in cyan in the other eye and vice versa for the object (Fig. 3 supplement 1). If cells followed color or eye-of-origin, then all the dots in the upper right quadrant in Fig. 3 supplement 1a should move to the lower left corner in Fig. 3 supplement 1b. Instead, the majority of cells followed the object identity rather than color or eye-of-origin for both the physical and perceptual condition (*p* < 10^−29^ for physical condition and *p* < 10^−11^ for perceptual condition, one-sided t-test, alternative hypothesis that modulation indices are greater than 0). This confirms that cells in IT cortex indeed represent the conscious percept rather than the color or eye-of-origin of the fixation spot.

To determine if one can decode the percept on a given trial from population activity, we performed recordings from multiple neurons simultaneously using S-probes with 32 electrode sites and passive Neuropixels-like probes with 128 electrode sites (see Methods for details). Fig. 4 shows recordings from face patch ML in one session using the Neuropixels probe. In this session, we recorded 81 cells simultaneously, of which 63 were face-selective (Fig. 4a). An example population time course snippet of cells recorded simultaneously in the perceptual switch condition showed clearly stronger activity across the recorded population during perception of face compared to object (Fig. 4b). The average population response across cells to perceptual switches is shown in Fig. 4c. We found above chance decoding of the perceptual condition in all 12 sessions (in all but one session, responses were recorded in both ML and AM, and cells were pooled across the two patches). Cross-validated accuracies of linear classifiers across different sessions are shown in Fig. 4d (see Methods). Decoding accuracies were 99% for the best session and 95% on average for the physical condition. For the perceptual condition, decoding was 88% on the best session and 78% on average.

**Figure 4.**
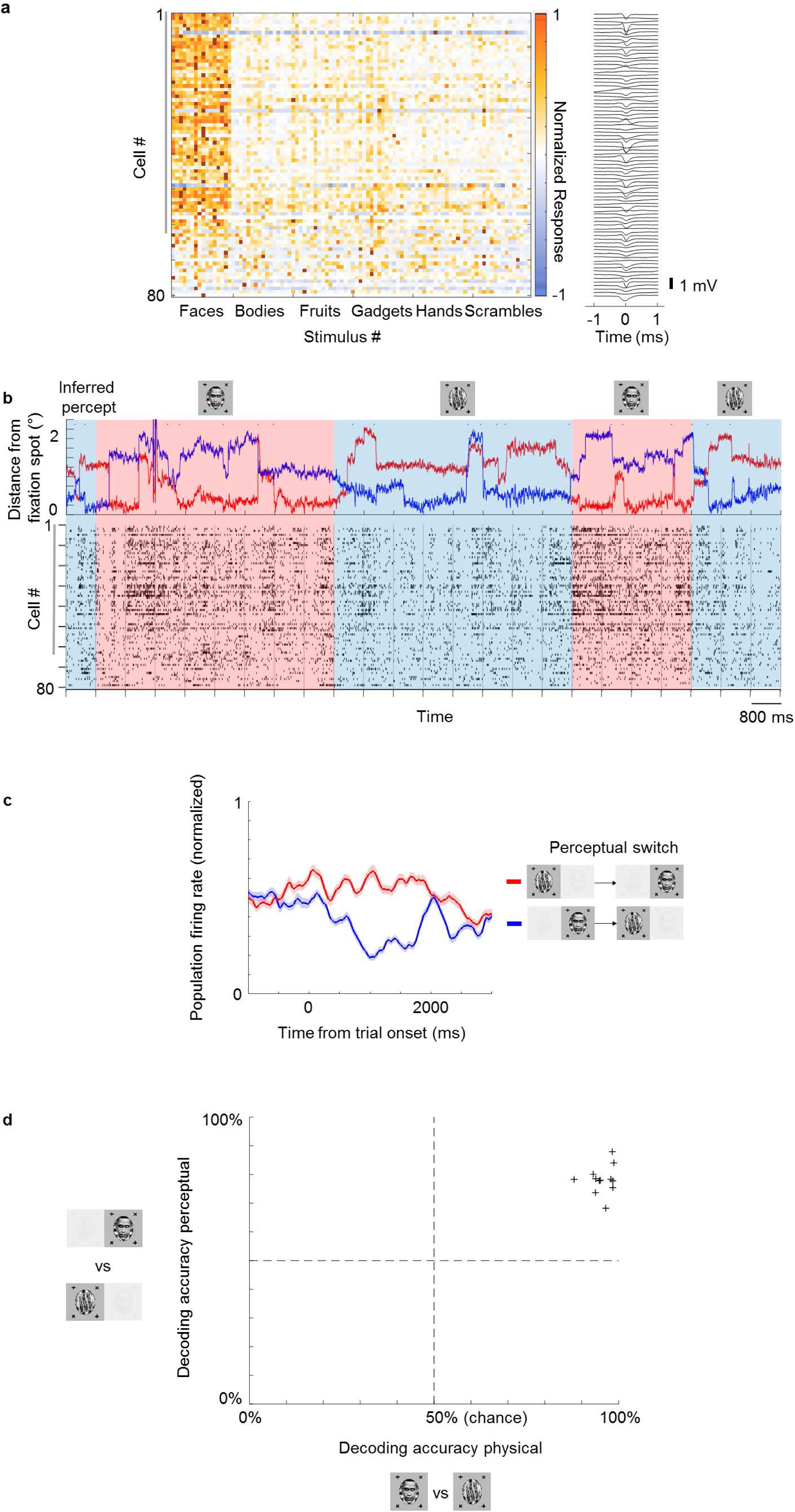
Multi-channel recordings allow decoding of conscious percept on single trials. **(a)** Left: Average responses (baseline-subtracted and normalized) of cells (rows) to 96 stimuli (columns) from 6 categories, including faces and other objects. Right: Waveforms of cells corresponding to rows on the left. Face-selective cells indicated by gray vertical bar on left. **(b)** Top: Example eye trace across 24 trials as in Fig. 1b in a binocular rivalry session (i.e. only perceptual, no physical switches). The inferred percept across trials according to eye trace is indicated by shading (red = face, blue = non-face object). Small black dots on top of eye traces indicate time points where our method detected saccades (see Methods), which were used in Fig. 5 and Fig. 5 supplement 1. Bottom: Response time course snippet of a population of 81 neurons recorded with a Neuropixels probe in ML simultaneously to the eye trace at top. Each row represents one cell; ordering same as in (a). Face-selective cells indicated by gray vertical bar on left. **(c)** Normalized average population response across all significantly face-selective ML cells recorded from one Neuropixels session (same session as in a, b) to perceptual switch from object to face (red) and face to object (blue). Shaded areas indicate standard error mean across cells. **(d)** Cross-validated decoding accuracy of a linear classifier trained to discriminate trials of inferred percept face versus inferred percept object for the physical switch condition (x-axis) and perceptual switch condition (y-axis). Each plus symbol represents a session of neurons recorded simultaneously with multi-channel electrodes.

Looking at the population time course, we noticed bursts of activity that appeared to be triggered by saccades, which occurred even when an object was perceived (blue epochs in Fig. 4b; small black dots on top indicate detected saccades). This raised the possibility that cells modulated by perception may still carry information about the physical stimulus. To investigate this further, we selected cells that (1) showed both significant physical and perceptual modulation and (2) consistently preferred the face over the object. We then averaged responses across these cells and computed response time courses triggered by individual saccades, grouped by whether a saccade occurred during a trial inferred to be face or object, respectively (Fig. 5). We observed response modulations for both physical and perceptual conditions starting around 130 ms after saccade onset (Fig. 5a). In the physical condition, a saccade during an object epoch led to response suppression, while a saccade during a face epoch led to response increase. In striking contrast, in the rivalry condition saccades led to response increase in both object and face epochs. As a consequence, during rivalry the response difference to a saccade between face and object, though significant (*p* = 10^−23^, two-sample t-test), was weaker than during the physical condition. Computing histograms of responses averaged across neurons for individual saccades shows that responses in the rivalry condition were less bimodal and spanned a smaller range compared to the physical condition (Fig. 5b). Importantly, this difference in response profiles between physical and perceptual conditions was apparent even when pooling across both face and object trials (Fig. 5b, middle), and *hence cannot be explained by mistakes in inferring the percept from eye movements*. We computed the absolute value of these responses and found the difference in response distributions to be significant (Figure 5b, right, *p* = 6 · 10^−35^, two-sample t-test on absolute value distributions).

**Figure 5.**
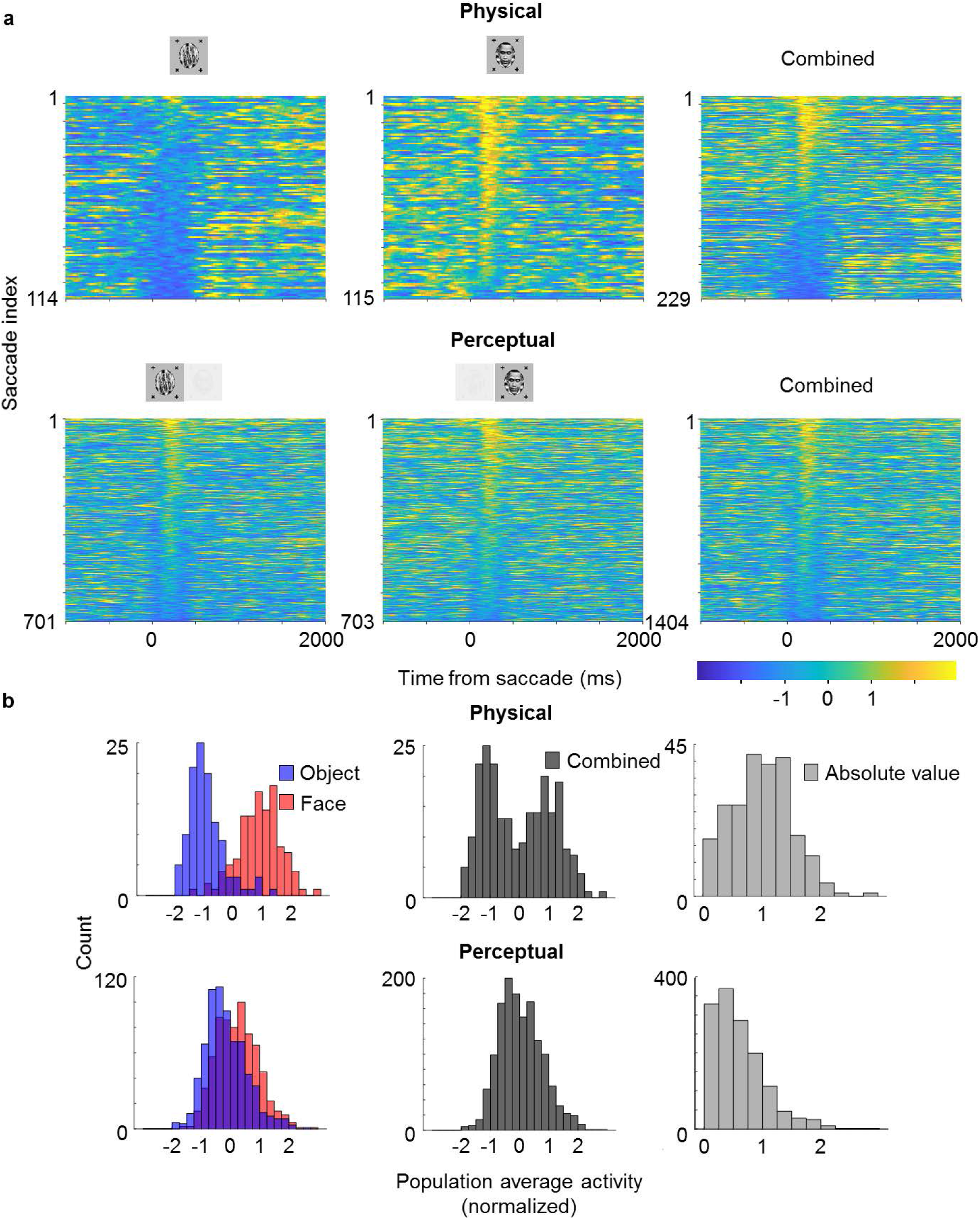
Saccade-triggered responses are less bimodal during rivalry. **(a)** Single-trial responses during saccades averaged across simultaneously recorded ML neurons from the same session as in Fig. 4b that were significantly face-selective for both physical and perceptual condition. Individual neuron responses were normalized to make -1 correspond to mean object response, 1 correspond to mean face response and 0 correspond to the average of the two. Rows of each plot correspond to response time courses to individual saccades, aligned to saccade onset, and sorted by average response during 0 to 400 ms after saccade onset. Top: Physical condition. Bottom: Perceptual condition. Left, middle, and right columns correspond to saccades during (inferred) object, face, and across both, respectively. The difference between perceptual and physical conditions in the third column shows that this difference cannot be simply attributed to mislabeling of perceptual state by the no-report paradigm. **(b)** Histograms of saccade-aligned responses averaged across a time window of 0 to 400 ms after saccade onset and across neurons (after normalizing as in (a)) that were significantly modulated for both physical and perceptual condition. Blue, red, and gray responses correspond to counts of saccade responses during object, face, and either, respectively. Top: Physical condition. Bottom: Perceptual condition. Left: Saccades for face and object plotted separately in red and blue, respectively. Responses were normalized to be 0 if the response was equal to the average of the face and object response, and 1 if equal to either the average face or average object response. Middle: Saccades for either face or object plotted in grey. Right: Absolute values of normalized responses plotted in light grey.

**Figure 3 supplement 1.**
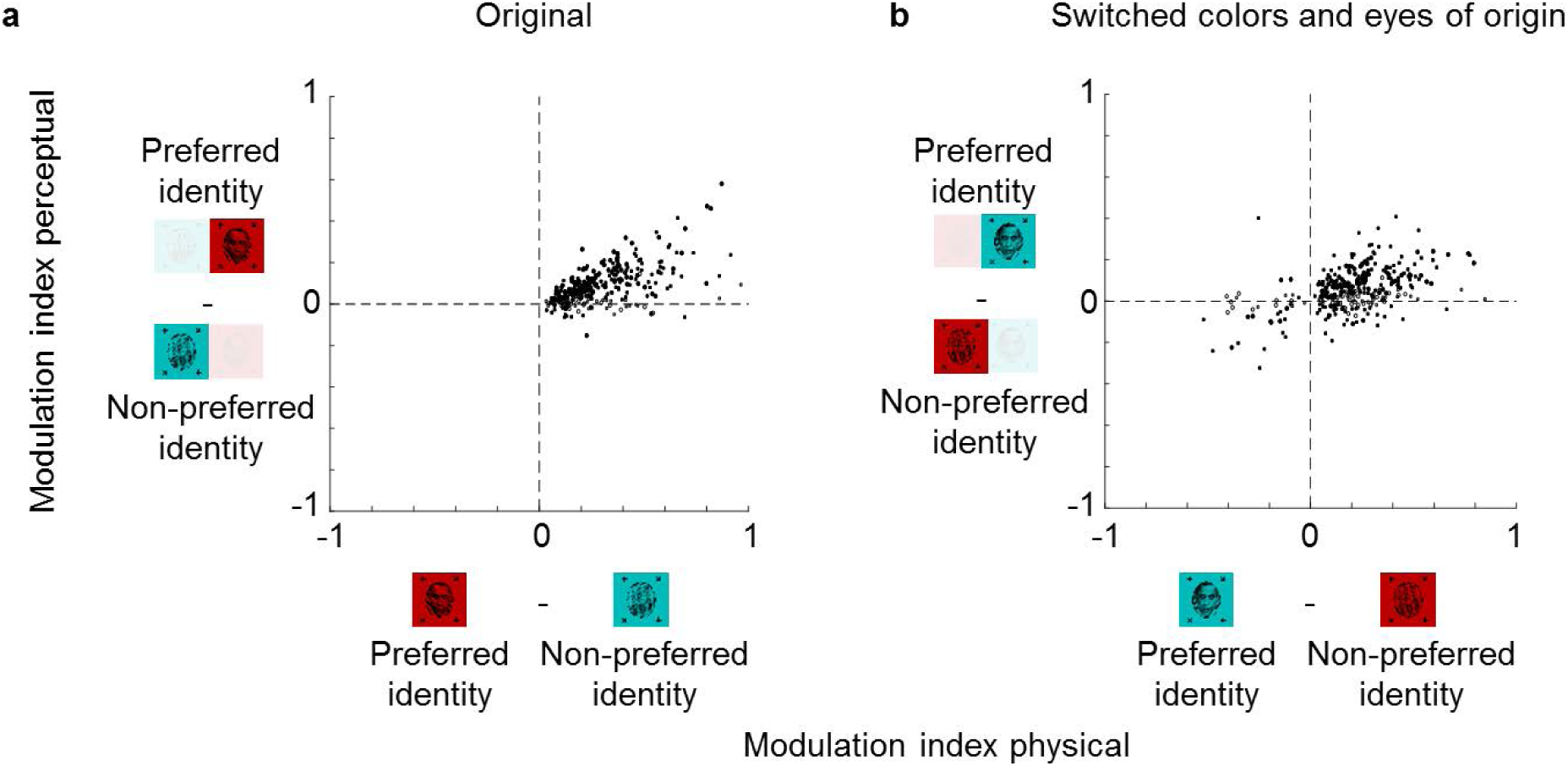
Color and eye-of-origin confound control. Left: Scatter plot similar to Fig. 3 but modulation indices 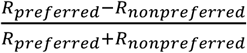 now show the difference between preferred and non-preferred stimulus. The preferred stimulus is face if the response to face is higher and non-face object if the response to non-face object is higher in the physical condition. Thus, by definition the x-values of all cells are positive. Right: Scatter plot of modulation indices 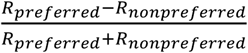 for the same preferred and non-preferred object identities of stimuli when the colors and eye of origin of the two stimuli were switched; importantly, the preference of a given stimulus identity was assigned based on responses to stimuli of the original color and eye of origin stimulus responses. N = 192 for ML and N = 120 for AM for both plots.

**Figure 5 supplement 1.**
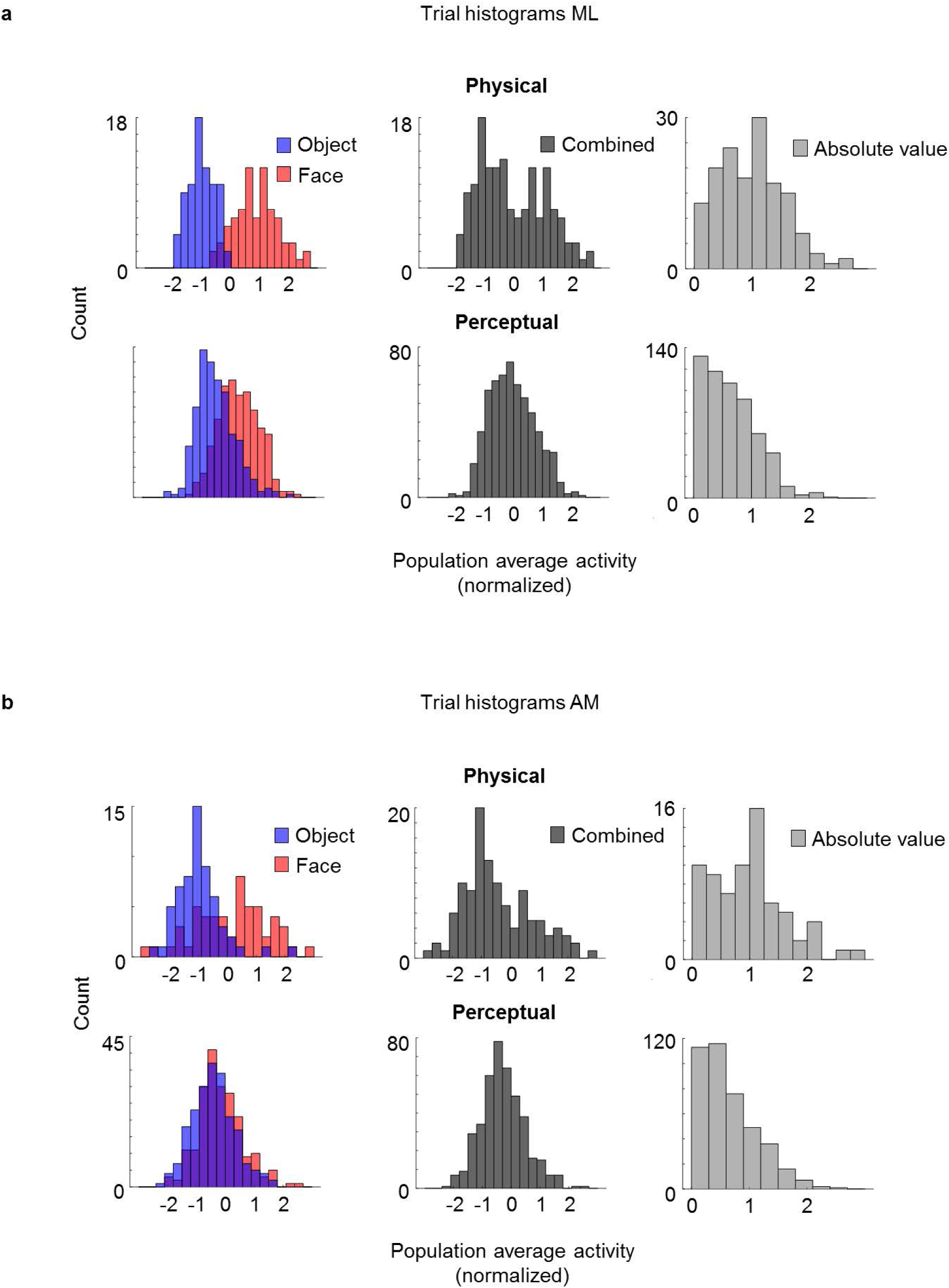
Lack of bimodality is a general trademark of rivalry. **(a)** Trial responses in ML are less bimodal during rivalry. Histograms have same conventions as Fig. 5b but instead of averaging neuron responses for individual saccades, responses are averaged across trial duration for individual trials. **(b)** Trial responses in AM are less bimodal during rivalry. Same conventions as in (a), but instead of the Neuropixels-like probe in ML, cells were simultaneously recorded from AM. Due to technical limitations, the 128-channels Neuropixels-like probe did not reach the depth of AM, and cells were recorded using a 32-channel S-probe instead.

The observation of different response profiles for physical and perceptual conditions was not specific for saccades: histograms were also less bimodal and spanned a smaller range for the rivalry condition when triggering responses on trial onsets rather than saccades in both ML (Fig. 5 supplement 1a, *p* = 9 · 10^−15^) and AM (Fig. 5 supplement 1b, *p* = 0.0014). Therefore, it appears that throughout rivalry, for perceptually-modulated cells, response differences between face and object are less pronounced than in the physical condition, and this is true in both ML and AM. One tantalizing explanation for this phenomenon is that perceptually-modulated cells may be multiplexing information about both the physical stimulus and the perceptual state during single trials, allowing both to be simultaneously represented across the face patch hierarchy.

## Discussion

We have shown that face patches ML and AM in macaque IT cortex are modulated by conscious perception and do not merely encode the physical input. Importantly, monkeys in this study had never been trained to actively report their percept. Instead, we were able to infer their percept from eye movements using a new no-report paradigm. Thus activity modulations attributed to switches in conscious perception in IT cannot be explained simply by active report.

Previous single-unit recordings in IT cortex using active report to infer the percept found 90% of cells represent the conscious percept (Sheinberg & Logothetis, 1997). Here, we found proportions of 61% in ML and 81% in the more anterior patch AM. The quantitative difference may be due to several factors including different recording sites (Sheinberg and Logothetis recorded from both upper and lower banks of the superior temporal sulcus in a less specifically targeted manner), imperfect accuracy of the no-report paradigm, and differences in stimuli and analysis methods. Importantly, our results show that the majority of cells in IT cortex do represent conscious perception and not merely active report and its accompanying cognitive factors. Furthermore, this new paradigm makes studies of consciousness in monkeys more accessible, by replacing the need to train the animal to signal its conscious percept (which can be a laborious process) with a simple task that only requires animals to follow a fixation spot.

Our results show that for cells that are modulated by conscious perception, the modulation is not “all-or-none.” First, we found that the average response modulation during the perceptual condition was weaker than during the physical condition (Fig. 3). This was also found in a previous study of rivalry (Sheinberg & Logothetis, 1997). This could be explained either by incomplete modulation, or by imperfect labeling of the animal’s perceptual state. The key question is: *what happens during single trials?* In the rivalry condition, do responses in single trials look like those to either physically-presented faces or objects? By recording from a large number of face cells simultaneously using a novel 128-electrode site probe specifically designed for use in primates, we could address this question for the first time. Surprisingly, we found a dramatically different response profile on single trials between the perceptual and physical conditions (Fig. 5). Whereas in the physical condition responses clustered into two groups, in the rivalry condition responses appeared unimodal, lying in between the two clusters for the physical condition. This suggests that single cells are multiplexing both the conscious percept and the veridical physical stimulus during single trials, such that information about both the perceived and unperceived stimuli remain constantly available in IT cortex. Future experiments varying the identity of the unperceived stimulus will be needed to further test this hypothesis. An alternative explanation is that cells are not modulated by the identity of the suppressed stimulus, and simply encode the dominant stimulus with reduced gain when presented in rivalry.

Compared to previous approaches that attempted to isolate representations of the conscious percept, our new no-report binocular rivalry paradigm has several advantages: For flash suppression, where a stimulus flashed in one eye suppresses the stimulus in the other eye, report is also not required (Tsuchiya & Koch, 2005; Wilke et al., 2003; Wolfe, 1984). However, in that case, the physical input when the target is perceived versus when it is suppressed are not identical, and thus any modulation observed may be driven entirely externally. Indeed, it is known that if a distractor stimulus is presented simultaneously with a preferred stimulus, the response can be reduced compared to when the preferred stimulus is presented alone as a result of simple normalization mechanisms (Bao & Tsao, 2018). Another paradigm that has been widely used to study the neural correlates of consciousness is backward masking. Here, the stimulus is presented for such a short time before being masked that sometimes it enters consciousness and sometimes not (Breitmeyer et al., 1984). So far, backward masking has always relied on report. Also, it is more susceptible to modulations arising from bottom-up withdrawal of attention or low-level (e.g., retinal) noise, whereas in binocular rivalry perceptual switches appear to be internally generated. One potential confound described by Block as the “bored monkey problem” is that the monkey may still be thinking about whether it is perceiving object or face and internally report it even if it is not required to actively report it (Block, 2020). It is methodologically very difficult to entirely remove this confound, but the fact that monkeys had to simultaneously perform a very challenging unrelated task of saccading to jumping fixation points should at least alleviate this concern. Thus, to the best of our knowledge, this study shows representations of the conscious percept in IT cortex in the most confound-free way to date. Our study complements a study conducted in parallel by Kapoor et al. (2020) that found modulations by conscious percept in prefrontal cortex using a different no-report paradigm based on optokinetic nystagmus.

The existence of two directly-connected functional modules with a hierarchical relationship (ML, AM) that both encode the conscious percept of a particular type of object opens up the possibility for future studies to investigate how changes in the conscious percept are coordinated across the brain. Recordings and perturbations in multiple face patches simultaneously using high-channel population recordings may reveal the dynamics of information flow, e.g., whether switches occur in a feedforward or feedback wave. This may yield insight into the mechanism for how a conscious percept emerges in the brain as an interpretation of the world that is consistent across different levels of representation.

## Methods

All animal procedures in this study complied with local and National Institute of Health guidelines including the US National Institutes of Health Guide for Care and Use of Laboratory Animals. All experiments were performed with the approval of the Caltech Institutional Animal Care and Use Committee (IACUC). The behavioral experiment with human subjects for the human psychophysics experiment complied with a protocol approved by the Caltech Institutional Review Board (IRB).

*Targeting*. Two male rhesus macaques were implanted with head posts and trained to fixate on a dot for juice reward. We targeted face patches ML and AM in IT cortex for electrophysiological recordings. ML and AM were identified using functional magnetic resonance imaging (fMRI). Monkeys were scanned in a 3T scanner (Siemens), as described previously (Tsao et al., 2006). MION contrast agent was injected to increase signal-to-noise ratio. During fMRI, monkeys passively viewed blocks of faces and blocks of other objects to identify face-selective patches in the brain. Recording chambers (Crist) were implanted over ML and AM. Guide tubes were inserted into the brain 4 mm past the dura through custom printed grids placed inside the chamber and electrodes were advanced to the target through the guide tube. Both chamber placement and grid design were planned with the software Planner (Ohayon & Tsao, 2012). After insertion of tungsten electrodes, correct targeting of the desired location was confirmed with anatomical MRI scans.

*Electrophysiology*. Recordings were performed using tungsten electrodes (FHC) with 1 MΩ impedance and, after correct targeting was confirmed, with 32-channel S-probes (Plexon) with 75 µm and 100 µm inter-electrode distance, and with passive Neuropixels-like probe prototypes (IMEC) (Dutta et al., 2019; Jun et al., 2017; Trautmann et al., 2019). These prototypes were a limited stock of test devices that were developed and used for testing as part of the development of primate Neuropixels probes and are not available for other labs. Unlike the final product, the prototypes had 128 passive electrode sites across 2 mm (arranged in two parallel staggered bands), but used the same electrode materials and shank specifications (45 mm total shank length). All electrodes were advanced to the target using an oil hydraulic Microdrive (Narishige). Neural signals were recorded using an Omniplex system (Plexon). Local field potentials were low-pass filtered at 200 Hz and recorded at 1000 Hz, and units were high-pass filtered at 300 Hz and recorded at 40 kHz. Only well-isolated units were considered for further analysis.

*Task*. Monkeys were head fixed and viewed an LCD screen (Acer) of 47 degree size in a dark room. Monkeys viewed stimuli of 5 degree size wearing red-cyan anaglyph goggles custom made with filters to match the red and green/blue emission spectrum of the screen, respectively, so that inputs to left and right eye could be controlled independently. Emission spectra were measured using a PR-650 SpectraScan colorimeter (Photo Research). Eye position was monitored using an eye tracking system (ISCAN). In the first phase of the experiment, monkeys passively viewed at least 5 repeats of 61 screening stimuli in pseudorandom order (250 ms ON time, 100 ms OFF time) with a fixation spot of 0.25 degree diameter in the center of the screen. Screening stimuli consisted of 20 images of faces and 41 images of non-face objects. During this phase, monkeys received a juice reward for maintaining fixation for at least 3 seconds. Subsequently, for the main experiment, stimuli contained one or two fixation spots at one of four possible locations (top, bottom, left, and right, 1 degree from the center) and were presented for 800 ms ON time and 0 ms OFF time. In the case of two fixation spots, stimuli contained one fixation spot per eye and the two spots never appeared at the same location. During this phase, the monkey received a juice reward if it maintained fixation within 0.5 degree of one of the fixation spot for at least half of the trial duration (i.e. 400 ms, not required to be contiguous). Stimuli during the main experiment included (1) a monocular face/monocular object with one fixation spot, and (2) a binocular stimulus composed of a face and a fixation spot in one eye, and an object and a second fixation spot in the other eye. To improve rivalry and reduce periods of mixture, face and object stimuli were presented on backgrounds consisting of gratings that were orthogonal in the two eyes. Moreover, we applied orthogonal orientation filters (with concentration *a*_*angle*_ = 0.5°) to the face and object stimuli, respectively, to increase local orientation contrast.

*Online analysis*. Spikes were isolated and sorted online using the PlexControl software (Plexon). During the screening phase, the average number of spikes during the time window from 100 ms to 300 ms was calculated for each unit and stimulus. For each stimulus, the average response across units was determined after normalizing the response of each unit by subtracting the mean and dividing by the standard deviation for the unit. Subsequently, the face stimulus with the highest average response and the object stimulus with the lowest average response were chosen to generate stimuli for the main experiment.

*Offline analysis*. For human subjects, the inferred percept based on button-presses on a given trial was determined according to the last report the subject made before the end of the trial. For humans and monkeys, we also determined their inferred percept based on eye movements depending on which fixation spot they fixated on if they fixated on one of the fixation spots for at least half of the trial duration (i.e. 400 ms, not required to be contiguous). We computed L-1 norms for computing the distance between eye position and a given fixation spot. We accounted for a saccade delay of on average 350 ms, by analyzing the eye data 350 ms until 1150 ms after trial onset For Figures 3 and supplement, 4d and 5 supplement, in order to exclude trials during which the percept switched back to the opposite percept, we also required the following trial to have the same inferred percept as the current trial. Spikes were re-sorted using the software OfflineSorter (Plexon). For Neuropixels, since the high density of electrodes allowed the same neuron to appear on multiple channels, we used Kilosort2 to re-sort spikes (Pachitariu et al., 2016). A total of 551 and 408 cells were recorded in monkey A and monkey O, respectively. To correct for delays in stimulus presentation, we used a photodiode that detected the onset and offset of the stimuli. The output of the photodiode was fed into the recording system and later used to synchronize the onset of the stimulus and the neurophysiological data during offline analysis. Peristimulus time histograms (PSTHs) were smoothed with a box kernel (100 ms width). For computing modulation indices we used the average spike count across trials as response. Decoding analysis was performed with a support vector machine with a linear kernel (Matlab fitcsvm) trained to discriminate trials where the inferred percept was face or object, respectively. As predictor variables we used the spike count during the 800 ms of each trial for all simultaneously recorded neurons. All decoding accuracies were cross-validated (leave-one-out). In more detail, one trial was chosen for testing and the rest of the trials for training and this was repeated for all trials to compute decoding accuracies. Criteria for detecting a saccade were as follows: A saccade was detected at time t if the distance between the mean eye position during t-100,…t-2 ms and the mean eye position during t+2,…t+100 ms was greater than 0.5 degree, and the eye position during t-100,…t-2 ms and t+2,…t+100 ms, respectively, stayed within 0.5 degree of the respective mean for at least 80% of the duration of each period. We also required consecutive saccades to be at least 100 ms apart from each other. All analysis was performed using Matlab (MathWorks).

## Author contributions

J.K.H. conducted the experiments and analyzed the data. J.K.H. and D.Y.T. designed the experiments, interpreted the data, and wrote the paper.

## Acknowledgements

This work was supported by HHMI and the Simons Foundation. We are grateful to members of the Tsao lab for feedback on the manuscript, Varun Wadia for helping us collect the human subject data, Audo Flores for animal support, Daniel Wagenaar and Eric Trautmann for technical assistance, and Barun Dutta, Tim Harris, Tirin Moore, Michael Shadlen, Krishna Shenoy, and HHMI for contributions to development of NHP Neuropixels probes.

